# A Modular Fusion Neural Network Approach to Efficiently Predict Multi-Metal Binding Sites in Protein Sequences

**DOI:** 10.1101/2025.09.13.676010

**Authors:** Jizheng Li, Changxin Fan, Hoi Ying Lau, Tianyue Wu

## Abstract

Accurate identification of metal-binding residues is essential for the study of metalloproteins, like zinc finger proteins, haemoglobin, and DNA polymerase. Due to the high cost and time demands of experimental methods, computational prediction has been applied. However, computational complexity and inextensibility of rigid frameworks impeded the application. This work presents a two-stage, sequence-based deep learning framework, which can predict zinc, iron, and magnesium binding amino acids of proteins. In the first stage, tokenized sequences are processed by independent one-dimensional convolutional neural networks (CNNs) to generate single-residue probability maps. In the second stage, a lightweight fusion network integrates these maps to model inter-metal dependencies and refine predictions. The framework employs an imbalance-aware loss function and the ensemble evaluation to improve robustness. Structural agnostic and modular design enable efficient training and inference, making it suitable for the annotation of large-scale proteome.

**CCS CONCEPTS:** • Applied computing; • Life and medical sciences; • Bioinformatics

## 1 INTRODUCTION

Metals play indispensable roles in molecular biology. As enzymatic catalytic centers, redox partners, and structural stabilizers, without metal ions such as zinc (Zn), iron (Fe), and magnesium (Mg), chemical transformations and molecular assemblies cannot be achieved. For example, Fe–S clusters shuttle electrons in respiratory chains [1], Zn^2+^ serves as a Lewis acid in hydrolytic enzymes and maintains the structural integrity of DNA-binding zinc fingers [2,3], and Mg^2+^ activates phosphoryl transfer reactions which also stabilizes RNA tertiary folds [4, 5]. Apart from catalysis and structural stabilization, metal ions also function in signal transduction [6], modulating the activity of ion channels, kinases, and transcription factors to orchestrate cellular responses to environmental and developmental cues [7].

Although there is an accumulation of high-resolution structures of metalloproteins in the Protein Data Bank, experimental identification of metal–protein interactions remains laborious, low-throughput, and costly [8, 9]. Traditional approaches including-ray crystallography, NMR spectroscopy, mass-spectrometry–based metalloproteomics, and affinity chromatography, require extensive samples under native conditions to preserve labile metal–protein complexes. These limitations hinder large-scale functional annotation of metal-binding sites and decelerate both fundamental research and drug discovery of metalloproteins.

To overcome these challenges, computational prediction of residue-level metal-binding sites has emerged as a powerful alternative. However, existing approaches face substantial computational and architectural limitations that constrain their practical applicability. Existing sequence-based multi-metal predictors often demonstrate suboptimal performance, or rely on rigid, metal-specific architectures that are tailored to individual ions, necessitating complete model redesign to accommodate additional metal types [10, 11]. Conversely, structure-based predictors, while potentially more accurate, require high-resolution 3D structures as input and rely on computationally expensive procedures such as homology modeling or structural alignment [12, 13, 14]. Recent approaches leveraging protein language models (e.g. ESM-1b, ESM-2) have shown promising accuracy but also demand substantial computational resources, with inference times significantly exceeding sequence-based methods and requiring extensive GPU memory for processing large amount of protein sequences [15, 16].

Here, we present a fusion neural network framework for multi-metal binding sites prediction that operates exclusively on primary protein sequence data. Our approach consists of two stages: (1) independent CNNs trained separately for Zn, Fe, and Mg to generate single-residue probability maps, and (2) a lightweight fusion network that integrates these maps to model inter-metal relationships and refine predictions. By integrating an imbalance-aware loss function, the ensemble evaluation, and a modular architecture, our model effectively addresses both the class imbalance between positive and negative samples for each metal and the complex inter-metal interactions within a unified framework, while achieving robust overall predictive performance. This structure-agnostic design enables rapid training and inference on large datasets without the requirement of structural inputs, which makes it ideally suited for the annotation of high-throughput proteome and a large-scale deployment.

## 2 MATERIALS AND METHODS

### 2.1 Dataset

A comprehensive dataset of metal binding proteins was retrieved from MbPA [17], from which proteins binding to Zn, Fe and Mg were selected for further analysis. The dataset comprises a total of 91,593 proteins, each of which is annotated with verified binding sites and their corresponding metal ions. The subset that can bind various metal ions is detailed in Table 1.

**Table 1:** Dataset.

|  | Zn | Fe | Mg |
| --- | --- | --- | --- |
| Binding proteins | 29424 | 26793 | 35374 |
| Binding residues | 115748 | 88795 | 81755 |

### 2.2 Sequence Encoding and Dataset Splitting Strategy

To establish a robust dataset for training and evaluation, all 91,593 protein sequences were firstly character-level tokenized and integer-encoded. Sequences shorter than 500 amino acids were post-padded with zeros at the C-terminus, while sequences longer than 500 amino acids were post-truncated from the C-terminus, yielding a uniform length of 500 amino acids. Simultaneously, a multi-label target tensor *Y* ∈ {0,1}^*N*×500×3^ (channels: Zn, Fe, Mg) was constructed by marking annotated binding-site positions for each metal, with all other entries set to zero.

To ensure balanced and reproducible evaluation, a per-sample stratification label was derived by aggregating each metal channel (logical OR) and the metal class with the highest presence was selected. Through these labels, 15% of the data was reserved as a stratified test set and the remaining 85% was used for model development. This development set was further split via stratified 5-fold cross-validation to facilitate hyperparameter tuning, early stopping, and unbiased final performance assessment.

### 2.3 Per-Metal Label Generation and Class Imbalance Handling

To enable independent training of metal-specific predictors while addressing class imbalance, a three-stage preprocessing pipeline was implemented:

#### 2.3.1 Per-Metal Label Generation

For each metal type *m* ∈ {Zn,Fe,Mg}, a dedicated binary label tensor *Y*^(*m*)^ ∈ {0,1}^*N*×500×1^ was constructed. Given annotated binding-site positions 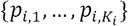 and their corresponding metal labels 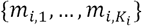 for sequence *i*, we set:

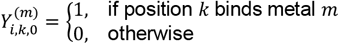

This yields metal-specific supervision masks essential for independent model training.

#### 2.3.2 Positive Sample Counting

To quantify class imbalance, positive labels per metal across the training set were counted: {*N*_Zn_, *N*_Fe_, *N*_Mg_}. These counts inform the subsequent loss weighting strategy.

#### 2.3.3 Weighted Binary Cross-Entropy Loss

Given the imbalance of the positive and negative binding labels, a custom loss function was defined:

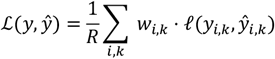

where *ℓ* is point-wise binary cross-entropy, *R* is the total number of residue positions, and the weight was defined as:

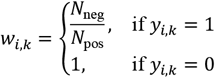

This amplifies penalties on misclassified binding residues, effectively mitigating class imbalance.

### 2.4 Single-Metal CNNs Architecture

Figure 1 illustrates the overall workflow of our model. A one-dimensional CNNs was employed to each single-metal model to predict position-wise binding probabilities for a specific metal ion. The architecture comprises:

**Figure 1:**
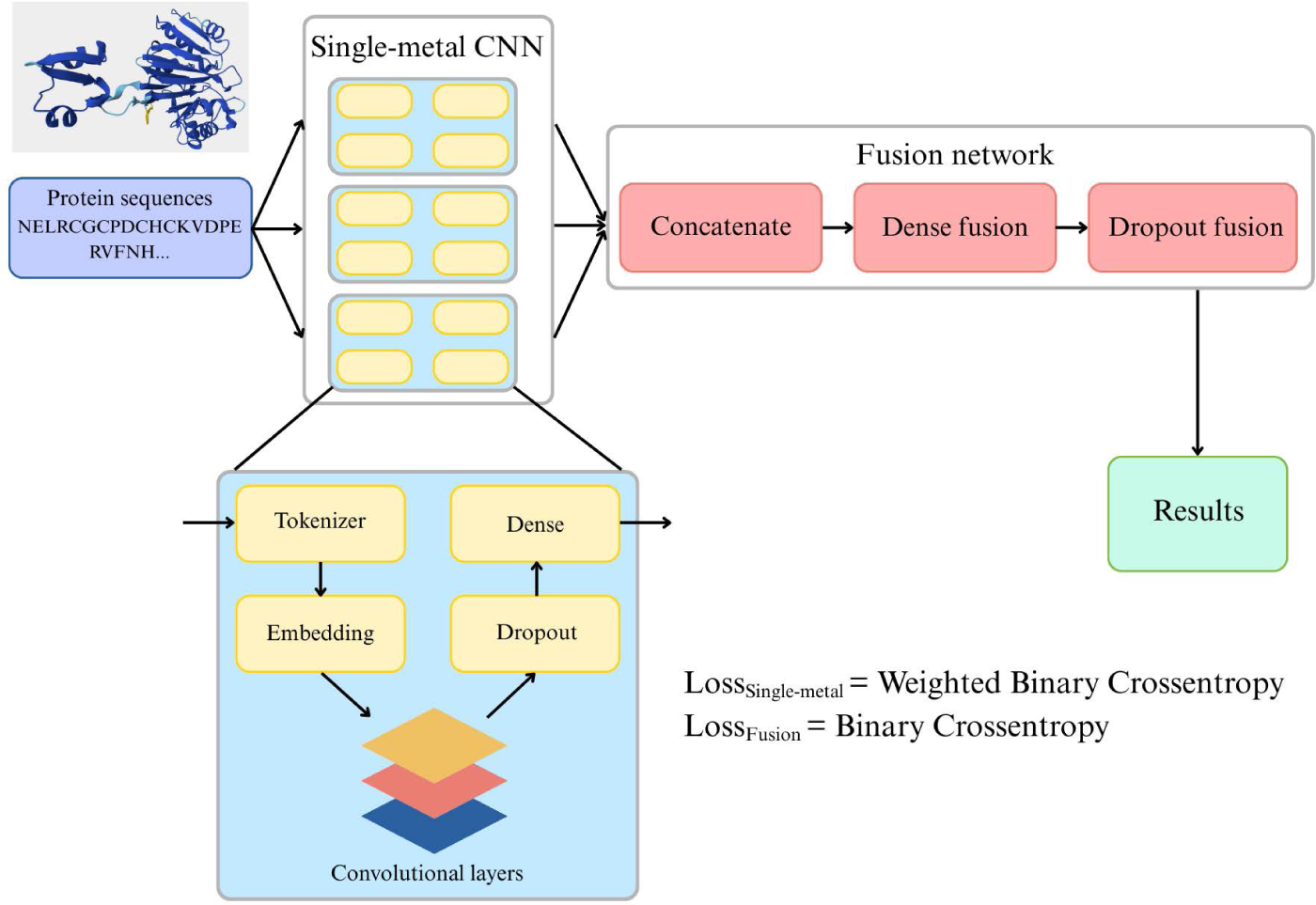
Overall framework of our fusion model.

- An embedding layer that maps each integer-encoded residue to a 64-dimensional trainable vector.
- Four Conv1D layers with filter {512,256,128,64} and kernel size {15,7,5,3} respectively, using ReLU activation
- A dropout layer (rate = 0.3) after the final convolution
- A Time Distributed Dense layer with sigmoid activation, yielding position-wise binding probabilities

Models were compiled with the Adam optimizer and the weighted loss function described above. Several hyperparameters were employed during the training process. The specific values of these hyperparameters are listed in Table 2.

**Table 2:** Training hyperparameters.

| Hyperparameters | Single-metal CNNs | Fusion network |
| --- | --- | --- |
| Epochs | 20 | 30 |
| Batch size | 128 | 128 |
| Patience | 3 | 5 |

### 2.5 Fusion Network for Multi-Metal Integration

To jointly predict binding probabilities across all three metal types, a lightweight fusion network was designed to ingest the position-wise outputs of the three single-metal models. For each sample, the concatenated predictions formed a tensor of shape (*L*_max_, *M*), where *L*_max_ = 500 amino acids and *M* = 3 metal channels. This tensor was passed through:

- A fully connected layer with 256 hidden units and ReLU activation, applied identically to each residue position to learn nonlinear interactions among metal-specific features.
- A dropout layer (rate = 0.2) to regularize the fusion weights and prevent overfitting.
- A final dense layer with *M* sigmoid outputs, yielding refined binding probabilities for Zn, Fe, and Mg at each residue.

The fusion model was compiled using standard binary cross-entropy loss—appropriate for the multi-channel, per-position binary classification task—and the Adam optimizer. The specific hyperparameters used for training the network is shown in Table 2. This fusion stage learned to correct correlated errors and leverage complementary strengths of individual metal predictors, improving overall binding site accuracy.

### 2.6 Performance evaluation

Let TP, TN, FP, and FN denote the total counts of true positives, true negatives, false positives, and false negatives aggregated over all residue–metal pairs in the test set. The following metrics are reported:

Precision

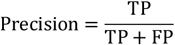

measures the fraction of predicted binding sites that are correct.

Recall

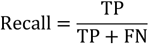

(also known as sensitivity) measures the fraction of true binding sites that are recovered.

F1 Score

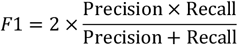

is the harmonic mean of precision and recall, balancing both aspects into a single measure.

Matthews Correlation Coefficient (MCC)

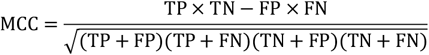

provides a balanced measure of binary classification quality even under class imbalance, with values in [−1,1].

These metrics collectively characterize the performances of the proposed model, ensuring a comprehensive evaluation of binding-site predictions.

## 3 RESULTS AND DISCUSSION

### 3.1 Structure-Agnostic Efficiency

Our framework is distinguished by its exclusive reliance on primary protein sequence data, thereby eliminating the computationally intensive requirements of homology modeling or structural alignment that are typically necessary for structure-based predictors. By encoding residue sequences as integer tokens and processing them through an embedding layer followed by CNNs, the entire training pipeline—including individual metal-specific submodels and the subsequent fusion network—can be completed in under one hour on a single NVIDIA A800 GPU. This efficiency facilitates rapid experimentation and real-time parameter tuning, which are critical for the high-throughput proteome annotation.

### 3.2 Threshold-Dependent Analyses

To validate the proposed fusion neural network, we performed stratified five-fold cross-validation on the development set to train robust models. Each model was subsequently evaluated on an independent test set, with performance metrics averaged across the five folds to ensure a reliable assessment of classification performance. For binary classification, a decision threshold τ was applied to the predicted binding probabilities: a residue is classified as a metal-binding site if its predicted probability exceeds τ; otherwise, it is designated as non-binding.

#### 3.2.1 F1-Score and MCC versus Threshold

Figure 2a illustrates the per-metal and macro-averaged F1 scores as functions of the decision threshold τ. As each thresholded prediction is derived from the mean output of the five fusion networks, the curves represent the ensemble’s smoothed probability distribution. The ensemble demonstrates robust performance for Fe, with F1 scores exceeding 0.81 for τ values between 0.25 and 0.60. For Zn and Mg, F1 scores surpass 0.79 within τ ranges of 0.25–0.50 and 0.25–0.60, respectively. The macro-averaged F1 score peaks at 0.855 within τ = 0.40– 0.45. Beyond τ = 0.50, the F1 score declines sharply due to an increase in false negatives, highlighting the importance of selecting τ within the optimal range of 0.40–0.45 to balance precision and recall across all metals.

**Figure 2:**
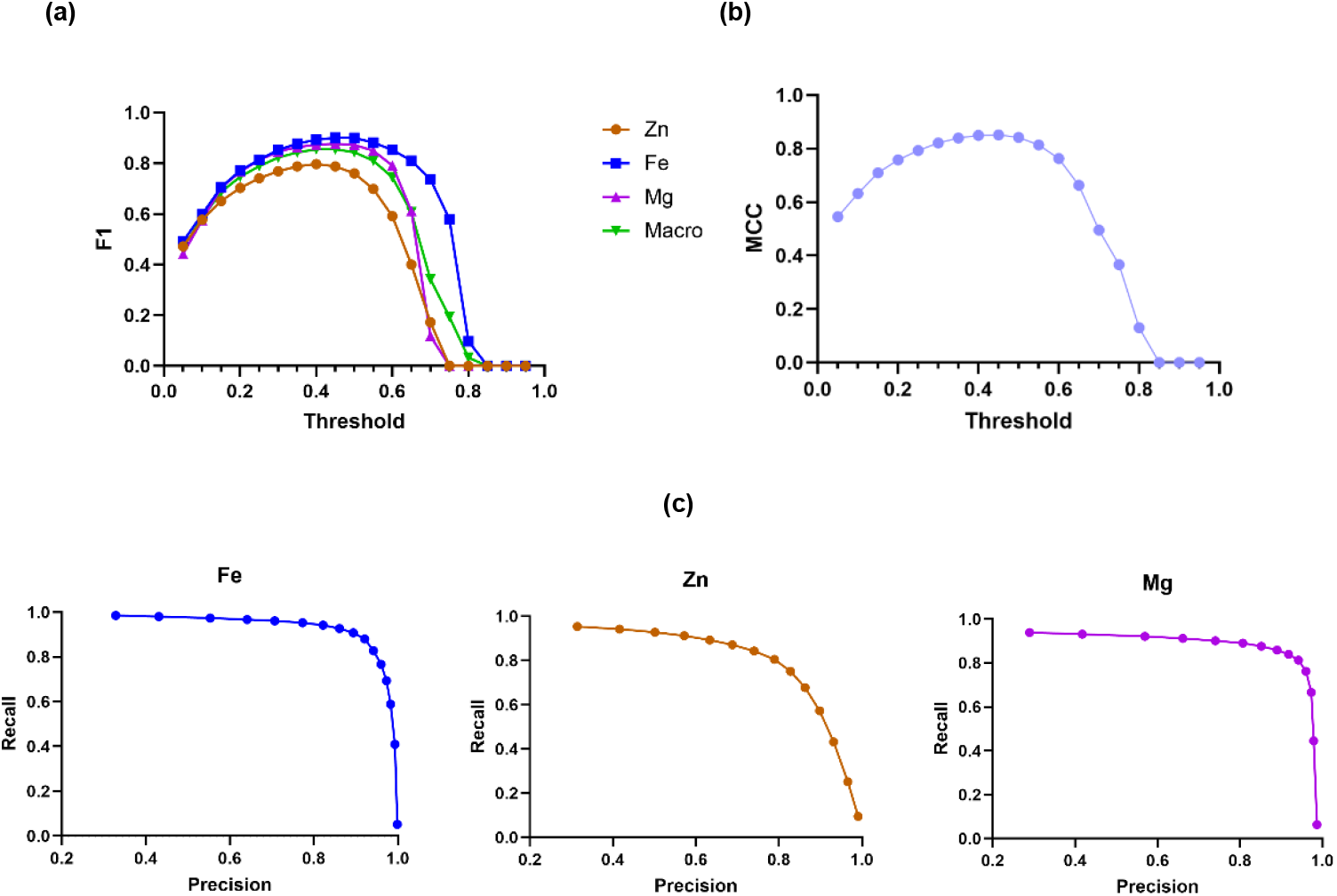
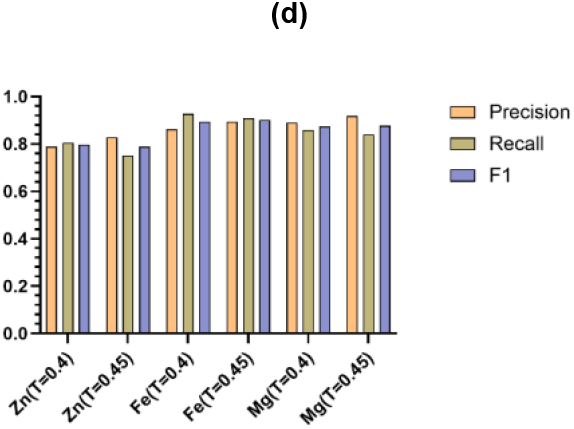
The performance evaluation results. (a) F1-Threshold curves of individual metal ions and their average, illustrating the performance of the macro fusion model. (b) MCC-Threshold curve of the macro fusion model. (c) Recall-Precision curves of three metal ions. (d) Model performance of different metal ions at thresholds of 0.4 and 0.45.

In Figure 2b, the MCC curve closely aligns with the macro-averaged F1 profile derived from the ensemble, further validating τ = 0.40–0.45 as the optimal operating range. This concordance confirms that the five-model ensemble achieves a balanced trade-off between true positive and false positive rates, even under conditions of severe class imbalance.

#### 3.2.2 Precision–Recall Trade-Off

Figure 2c presents precision–recall curves for each metal, derived from thresholded ensemble predictions. Predictions of Fe binding maintain high precision at elevated recall levels, making them suitable for comprehensive site screening. Predictions of Zn binding, despite a strong AUC, require conservative thresholding in applications prioritizing precision over coverage. Predictions of Mg binding exhibit precision above 0.80 up to a recall of approximately 0.90, with a gradual decline, indicating a balanced performance across thresholds. This makes the ensemble robust for applications requiring both moderate high recall and sustained precision.

### 3.3 Per-Metal Performance at the Optimal Threshold

At the selected decision thresholds, Figure 2d illustrates the ensembled model’s precision, recall, and F1 scores for each metal class. Fe achieves the highest F1 score by maintaining an optimal balance between precision and recall, making it ideal for tasks that require both high sensitivity and specificity. Mg follows with a strong F1 score, supported by high precision and recall, demonstrating consistent performance across various thresholds. In contrast, Zn exhibits high precision but lower recall at the higher threshold, which improves with a slight threshold adjustment, thereby enhancing its F1 score. These metal-specific trade-offs underscore the ensemble’s flexibility, allowing for the prioritization of precision for targeted experimental validation and the recall for comprehensive, high-throughput annotation.

Regarding the macro-averaged performance, as shown in Table 3, the ensembled model delivers robust results across all metal classes. It maintains high overall performance, with the optimal threshold range effectively balancing precision and recall, as evidenced by stable F1 scores and strong correlation metrics.

**Table 3:** Macro Fusion Model Performance Averaged Over 5-Fold Cross-Validation.

| Threshold | Precision | Recall | F1 | MCC | AUC | AUPRC |
| --- | --- | --- | --- | --- | --- | --- |
| $\tau = 0.4$ | 0.846 | 0.864 | 0.855 | 0.850 | 0.996 | 0.898 |
| $\tau = 0.45$ | 0.880 | 0.833 | 0.855 | 0.851 | 0.996 | 0.898 |

### 3.4 Ablation study

To systematically evaluate the contribution of each architectural component, we conducted a comprehensive ablation study using stratified 5-fold cross-validation on the development set. Table 4 presents the averaged performance metrics across all folds, demonstrating the progressive improvements achieved by each design choice.

**Table 4:**
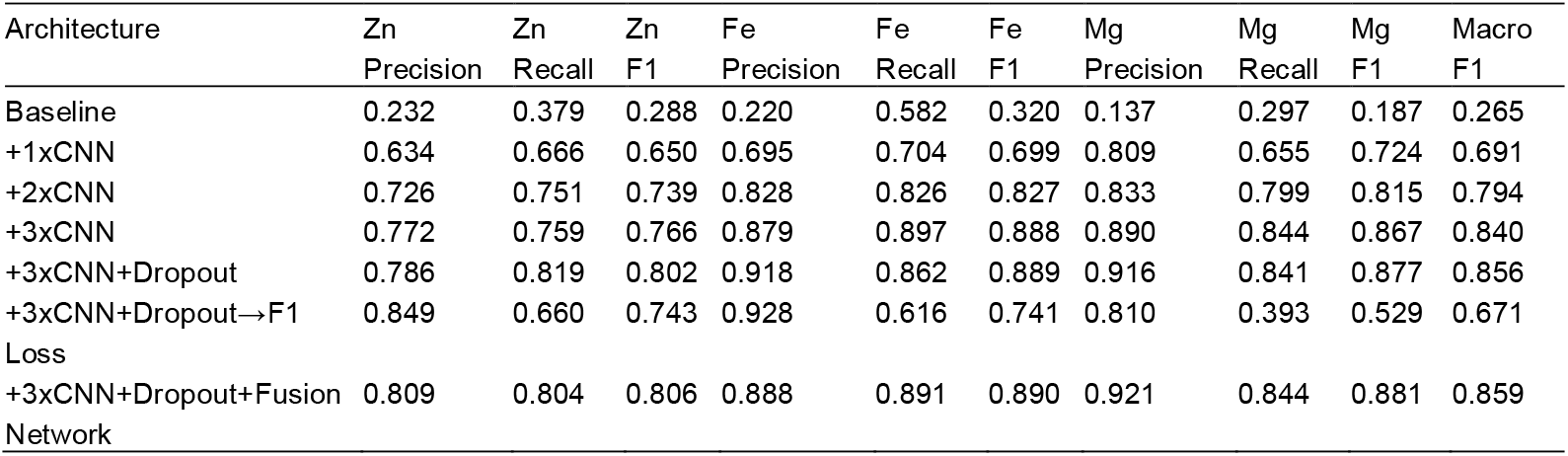
Results of the ablation study.

#### 3.4.1 CNNs and Dropout

The baseline model, which employed basic sequence encoding and a single CNN layer, initially exhibited suboptimal performance, achieving a macro-averaged F1 score of 0.265 across all metal types. The sequential incorporation of additional CNN layers significantly enhanced performance, with the F1 score rising to 0.840 (an increase of 0.575) after adding three CNN layers. This marked improvement highlights the critical role of hierarchical feature extraction, where deeper layers capture complex sequence motifs essential for accurate metal-binding site identification.

Moreover, incorporating a dropout layer after the third convolution raised macro F1 to 0.856 (+0.016), demonstrating its role in preventing overfitting and improving generalization without reducing sensitivity to rare binding sites.

#### 3.4.2 Imbalance-Aware Loss Function

Metal-binding sites exhibit class imbalance at the residue level, with negative (non-binding) positions vastly outnumbering positive (binding) annotations. To address this fundamental challenge, we implemented a weighted binary cross-entropy loss function that dynamically adjusts the penalty for misclassifying rare positive samples. As shown in Table 4, while the F1-based loss primarily emphasizes precision, our weighted loss substantially improves recall without compromising overall balance, thereby yielding superior F1 scores across all metal ions and in the macro-averaged evaluation. By increasing the gradient contribution from scarce binding residues during backpropagation, the weighted loss counteracts the model’s tendency to favor the dominant negative class. Consequently, it achieves higher recall and F1 for metal-binding sites without resorting to complex resampling methods, ensuring balanced learning across both abundant non-binding positions and rare binding residues.

#### 3.4.3 Fusion Layer

Integrating metal-specific CNNs outputs via a lightweight fusion layer produced a modest yet meaningful improvement in macro F1 (0.859). Calibration was significantly enhanced, reducing the precision–recall discrepancy from 0.056 to 0.003 for Fe and from 0.033 to 0.005 for Zn, thereby achieving more uniform predictive performance across metals. Such refined calibration permits the adoption of a single decision threshold, obviating the need for discrete threshold optimization for each ion and simplifying application to proteins with multiple binding metals. Furthermore, the fusion layer effectively models inter-metal dependencies: for Fe, precision and recall shifted from a precision-dominant 0.918/0.862 to a balanced 0.888/0.891. By leveraging co-occurrence patterns among metal-binding sites, the fusion network enhances both accuracy and robustness of residue-level predictions.

The ablation results collectively validated two core design principles: (1) the weighted binary cross-entropy loss function is essential for handling class imbalance in metal-binding site prediction, and (2) the fusion network architecture, while providing modest aggregate improvements, enhances prediction consistency and captures valuable cross-metal relationships that individual models cannot exploit independently.

## 4 CONCLUSION

This study presents a robust and extensible deep learning framework for predicting multi-metal binding sites in protein sequences. It utilizes a two-stage architecture that combines specialized CNNs with a fusion network for integrative analysis. By relying exclusively on primary sequence information and incorporating imbalance-aware loss functions, this approach enables rapid, high-throughput, and accurate annotation of Zn, Fe, and Mg binding sites without structural data or costly computational modeling.

Stratified dataset splitting and systematic cross-validation establish reliable performance benchmarks and foster reproducibility. Empirical results demonstrate that the fusion network effectively captures cross-metal interdependencies, achieving consistently high precision and F1 scores across diverse benchmarking scenarios. Moreover, the modular design of our pipeline enhances both scalability and extensibility. Each metal ion is modeled independently using dedicated one-dimensional CNNs that are trained exclusively on their respective binding labels. This modularity allows for the seamless integration of additional metal classes by simply training new submodels with an identical architecture and incorporating their output into the existing fusion network. Consequently, the framework implements a plug-and-play paradigm for multi-metal binding-site prediction that can be readily extended to other metals.

Importantly, the combination of targeted preprocessing, adaptive loss weighting, and ensemble-based evaluation directly addresses the challenge of class imbalance and enhances sensitivity for rare binding residues. The efficiency of the training pipeline supports rapid experimentation and large-scale deployment, making the framework suitable for proteome-wide studies and facilitating collaborative extension.

In summary, this fusion neural network approach advances metalloprotein annotation by offering a sequence-based, scalable, and high-performance solution, accelerating the discovery and analysis of biologically significant metal-protein interactions in both research and biomedical contexts.

## Acknowledgements

The authors express their sincere gratitude to all 2025 HKUST International Genetically Engineered Machine (iGEM) advisors and team members, with special thanks to Professor King Lau Chow and Dr. Jessica Ce Mun Tang for their invaluable guidance and support.

